# PathWalks: Identifying pathway communities using a disease-related map of integrated information

**DOI:** 10.1101/2020.01.27.921270

**Authors:** Evangelos Karatzas, Margarita Zachariou, Marilena Bourdakou, George Minadakis, Anastasios Oulas, George Kolios, Alex Delis, George M. Spyrou

## Abstract

Understanding disease underlying biological mechanisms and respective interactions remains an elusive, time consuming and costly task. The realization of computational methodologies that can propose pathway/mechanism communities and reveal respective relationships can be of great value as it can help expedite the process of identifying how perturbations in a single pathway can affect other pathways.

Random walks is a stochastic approach that can be used for both efficient discovery of strong connections and identification of communities formed in networks. The approach has grown in popularity as it efficiently exposes key network components and reveals strong interactions among genes, proteins, metabolites, pathways and drugs. Using random walks in biology, we need to overcome two key challenges: 1) construct disease-specific biological networks by integrating information from available data sources as they become available, and 2) provide guidance to the walker so as it can follow plausible trajectories that comply with inherent biological constraints.

In this work, we present a methodology called PathWalks, where a random walker crosses a pathway-to-pathway network under the guidance of a disease-related map. The latter is a gene network that we construct by integrating multi-source information regarding a specific disease. The most frequent trajectories highlight communities of pathways that are expected to be strongly related to the disease under study. We present maps for *Alzheimer’s Disease* and *Idiopathic Pulmonary Fibrosis* and we use them as case-studies for identifying pathway communities through the application of PathWalks.

In the case of *Alzheimer’s Disease*, the most visited pathways are the “Alzheimer’s disease” and the “Calcium signaling” pathways which have indeed the strongest association with *Alzheimer’s Disease*. Interestingly however, in the top-20 visited pathways we identify the “Kaposi sarcoma-associated herpesvirus infection” (HHV-8) and the “Human papillomavirus infection” (HPV) pathways suggesting that viruses may be involved in the development and progression of *Alzheimer’s*. Similarly, most of the highlighted pathways in *Idiopathic Pulmonary Fibrosis* are backed by the bibliography. We establish that “MAPK signaling” and “Cytokine-cytokine receptor interaction” pathways are the most visited. However, the “NOD receptor signaling” pathway is also in the top-40 edges. In *Idiopathic Pulmonary Fibrosis* samples, increased NOD receptor signaling has been associated with augmented concentrations of certain strains of Streptococcus. Additional experimental evidence is required however to further explore and ascertain the above indications.

## Introduction

Since its introduction more than a century ago, Random Walks [1] have been successfully applied to a wide range of sciences including Physics, Chemistry, Biology, Computer Science and Engineering. With its effective algorithmic layout, easy realization and efficiently produced outcomes, the method is still deemed a suitable choice for extracting sub-networks of interest in graph-structures consisting of nodes demonstrating multiple strong connections. The methodology has known weaknesses such as simply recreating the degree distribution of a graph or getting trapped in highly connected cliques without being able to explore distant neighborhoods. Moreover, random walks entail a finite number of steps and in this respect, if additional neighborhoods are to be explored during the same time period, multiple walkers have to be simultaneously deployed [2, 3]. To prevent walkers from entrapment in strongly-connected network regions, restart strategies are used [4, 5]. Such restart strategies allow a walker to discontinue its current walk and proceed by following up a different node in the graph. Converging strategies have also been studied mostly in the context of computer networks where random walks converge according to application-induced probability distributions for visiting nodes [6].

The output from methodologies such as the Random Walk is heavily dependent on the quality of the contained data. In random walks specifically, these data are integrated in a graph. There is a vast number of online databases containing biological content and an even greater need of parsing and integrating this information [7–9]. The potential knowledge gain could provide researchers with the means and tools to extract results that would benefit the health care system by enhancing prevention, diagnosis as well as treatment of maladies. Computational applications which allow for fast screening and integration of such biological information are the prerequisite for speeding up the process of generating quality results. In this respect, tools such as the PREDICT [10] integrate drug information from online databases including DrugBank [11], OMIM [12] and SIDER [13] in order to suggest new drug-target indications based on substance similarities. Other models including MutPred [14] parse protein sequences and provide insights on the mechanisms of diseases. Similar software tools are especially needed when it comes to rare diseases as *in vivo* experiments might not be given the appropriate consideration. The latter could be attributed to the lack of targeted individuals especially if a disease under examination is simply infrequent.

In the work of Zachariou et al. (2018) [15], the importance of studying disease mechanisms from a multi-omics perspective was examined and a multi-level network for the *Alzheimer’s Disease* (*AD*) was proposed. This network was formed by integrating multi-source biological information such as differentially expressed genes, pathways, single nucleotide polymorphisms, drugs and microRNAs. Here, genes act as intermediaries between the different layers of the proposed network. Through this methodology, clusters of potential key biological pathways of *AD* were proposed for further examination.

Community detection algorithms are regularly used in order to identify meaningful clusters in a graph and have been successfully proposed in the context of social networks for more than a decade now [16–18]. We have only recently seen the adoption of such techniques in biological settings. In particular, a benchmarking study [19] considered the Louvain method [20] as the best choice in finding protein communities in the protein-protein interaction (PPI) networks of Human and Yeast. While addressing the DREAM challenge, Tripathi et al. [21] applied their community detection framework in six heterogeneous biological networks (two human PPI, a pathway signaling, a co-expression, a cancer and a homology network) in order to extract core disease communities. More specifically, they showed that overlapping community detection algorithms yield better results for disease module identification, which is justified since a node (e.g. a gene) can participate in multiple diseases at the same time. Wilson et al. [22] applied community detection algorithms in a gene interaction network and while deploying the Louvain algorithm they sought to identify communities of up to ten genes that characterize functional and disease pathways.

In this work, we propose a random walk-based methodology on a pathway-to-pathway network and we term this as *PathWalks*. PathWalks exploits a map, that we construct in the form of a synthetic gene network, containing integrated information regarding a disease of interest, as the latter has been presented in [15]. We create multisource integrated information maps regarding *AD* and *Idiopathic Pulmonary Fibrosis* (*IPF*). We use the produced maps to drive random walks on a reference pathway-to-pathway network. Our methodology highlights the most frequently walked trajectories, identifying pathway communities that are expected to be strongly related to these diseases. The novelty of our PathWalks approach lies with the exploitation of multi-omics disease-related information that helps drive walks on a functional connectivity network of biological pathways. The approach ultimately highlights pathway communities related to a disease of interest.

## Materials and Methods

### The General Concept of PathWalks

Our proposed PathWalks methodology performs random walks on a pathway-to-pathway network under the guidance of a synthetic gene network that we construct by integrating a-priori molecular information related to a disease [15]. The PathWalks methodology makes use of two main network components which are related to a disease of interest and need to be constructed before the execution of the algorithm. The first component is the multisource information map; this is a synthetic gene-to-gene network which represents integrated information (e.g. gene co-expression, physical interactions, miRNA targets) from biological databases in the form of connections and edge weights. Mathematically, the gene network is represented as a graph (*G_g_)* and described as *G_g_ = (V_g_, E_g_)*, where *V_g_* is the set of nodes (genes) and *E_g_* is the set of connections between the nodes of the graph. The walker performs random walks on the gene network and the visited nodes indicate the walker’s destination on the PathWalks’ second component; the biological pathways’ network.

We construct the pathway-to-pathway network (*G_p_ = (V_p_, E_p_*), by parsing the biological pathways’ functional connectivity information existing in KEGG [23]. Pathways that contain genes already associated with the studied disease, receive higher numeric-value edge scores (i.e. visitation probability). The walker moves on the pathway-to-pathway network according to the instructions given by the map (gene-to-gene network) in order to explore biological pathway relations regarding the disease under examination. A sorted list of the most visited pathways is generated after a set number of iterations. In order for the algorithm to converge, the two last sorted pathway-visitation lists must have a similarity index above a selected threshold. Finally, the algorithm highlights the most frequently visited edges (i.e. pathway-to-pathway connections) and nodes (pathways), revealing interesting pathway communities, according to the multisource map. In this study, we explore two use case scenarios from different disease settings; *AD* as a neurodegenerative disease and *IPF* as a fibrotic disease. We show a descriptive diagram of the PathWalks methodology in Figure 1.

**Figure 1:**
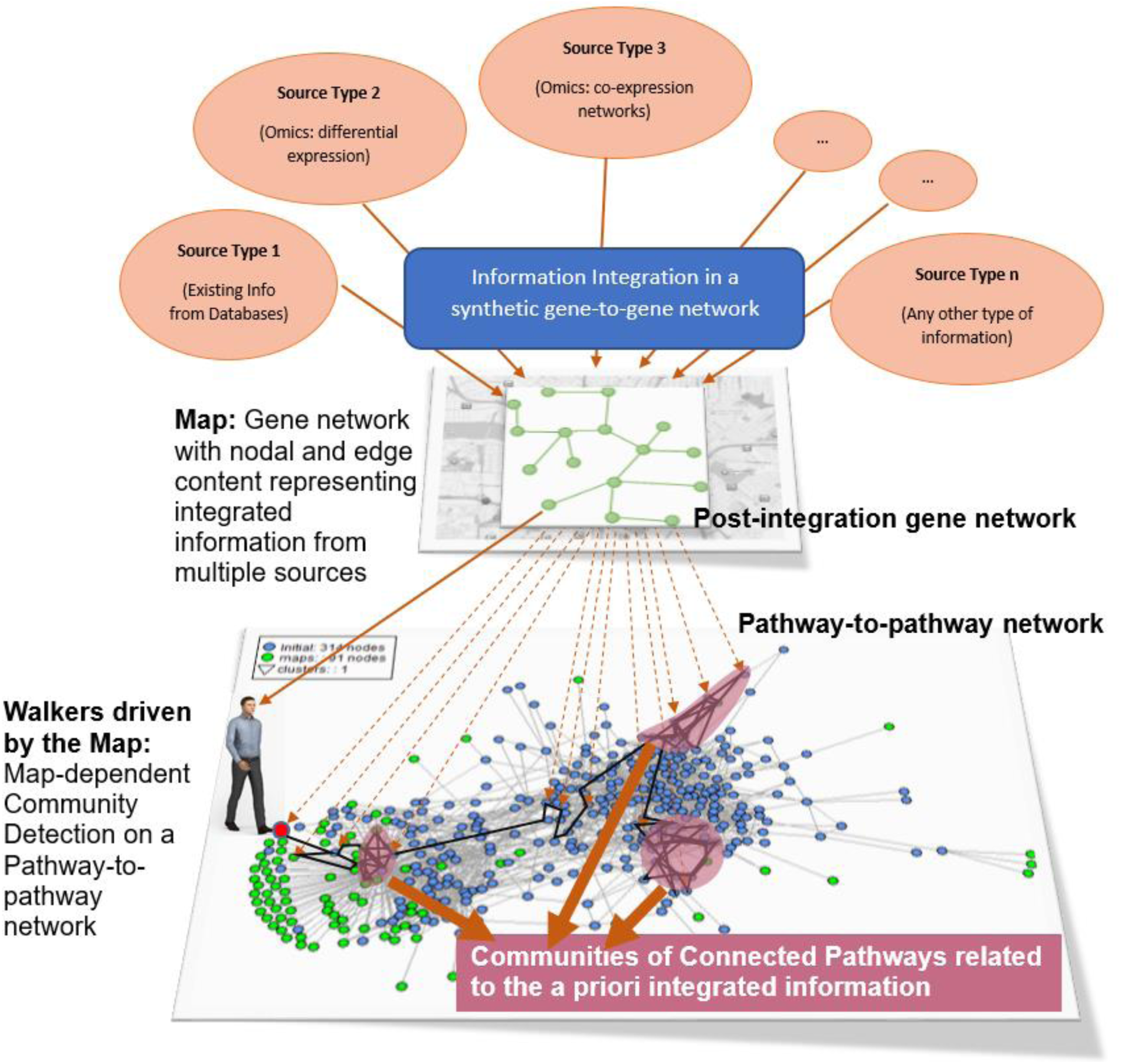
The PathWalks Concept. Multisource information regarding a disease is integrated in a gene map. This gene map guides the walker on a functional connectivity network of biological pathways in order to identify key pathway communities of the disease.

### Multisource Integrated Gene Map per Disease

The first component needed for the execution of PathWalks is the gene map. Here, we create gene maps for the PathWalks algorithm by integrating biological information as described [15]. For both *AD* and *IPF* maps, we downloaded genes, drugs, biological pathways and single nucleotide polymorphisms from Malacards [24]. For the *AD* map, we further included copy number variations’ information from Malacards, which was missing in the case of *IPF*. We linked drugs of both cases to their gene targets via the DrugBank database. We then extracted additional genetic and physical interaction information for each disease’s genes through GeneMANIA [25]. Finally, we mapped the genes of each disease to miRNAs through the MirTarBase [26] database. In the *AD* use case, additional miRNAs were explored through miRBase [27] and TargetScan [28]. Following the multisource integration, we prepared the generated gene-to-gene networks to be used as guiding maps in the PathWalks execution.

We visualize the gene maps for *AD* and *IPF* in Figures 2 and 3 respectively. In these figures the node size is relative to the node’s degree while the edge width is relative to the edge weight (gene-gene relation scores). A node’s degree represents its total number of connections. Intuitively, nodes with multiple high-score connections (larger node size and edge thickness) are more likely to be accessed by the walker, therefore they are the main candidates driving the walks on the pathways’ level.

**Figure 2:**
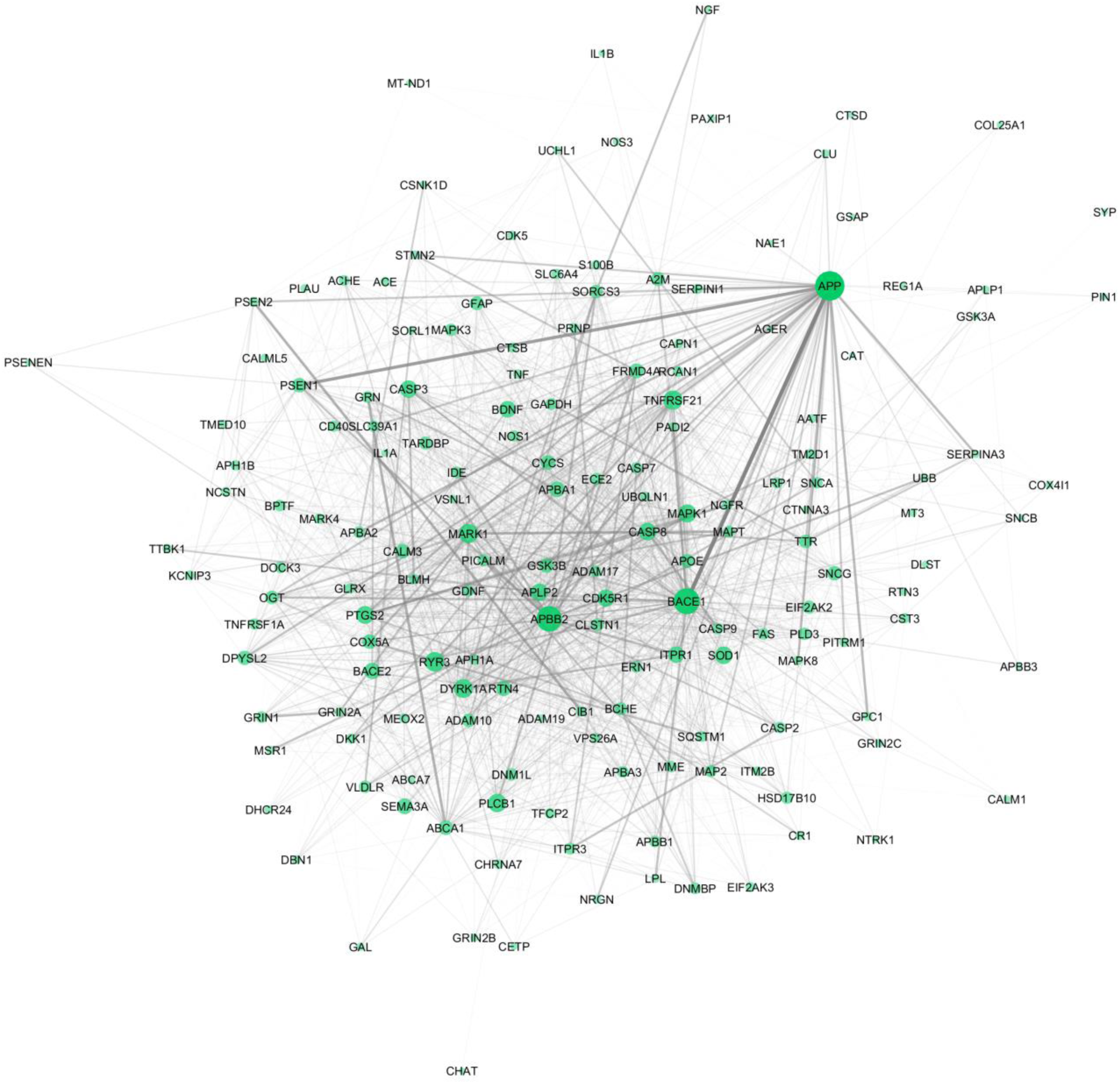
The gene map/network for the use case of AD. The network consists of 165 gene nodes and 1965 connections. The node size and transparency are based on the node’s degree and the edge width and transparency on the edge weights.

**Figure 3:**
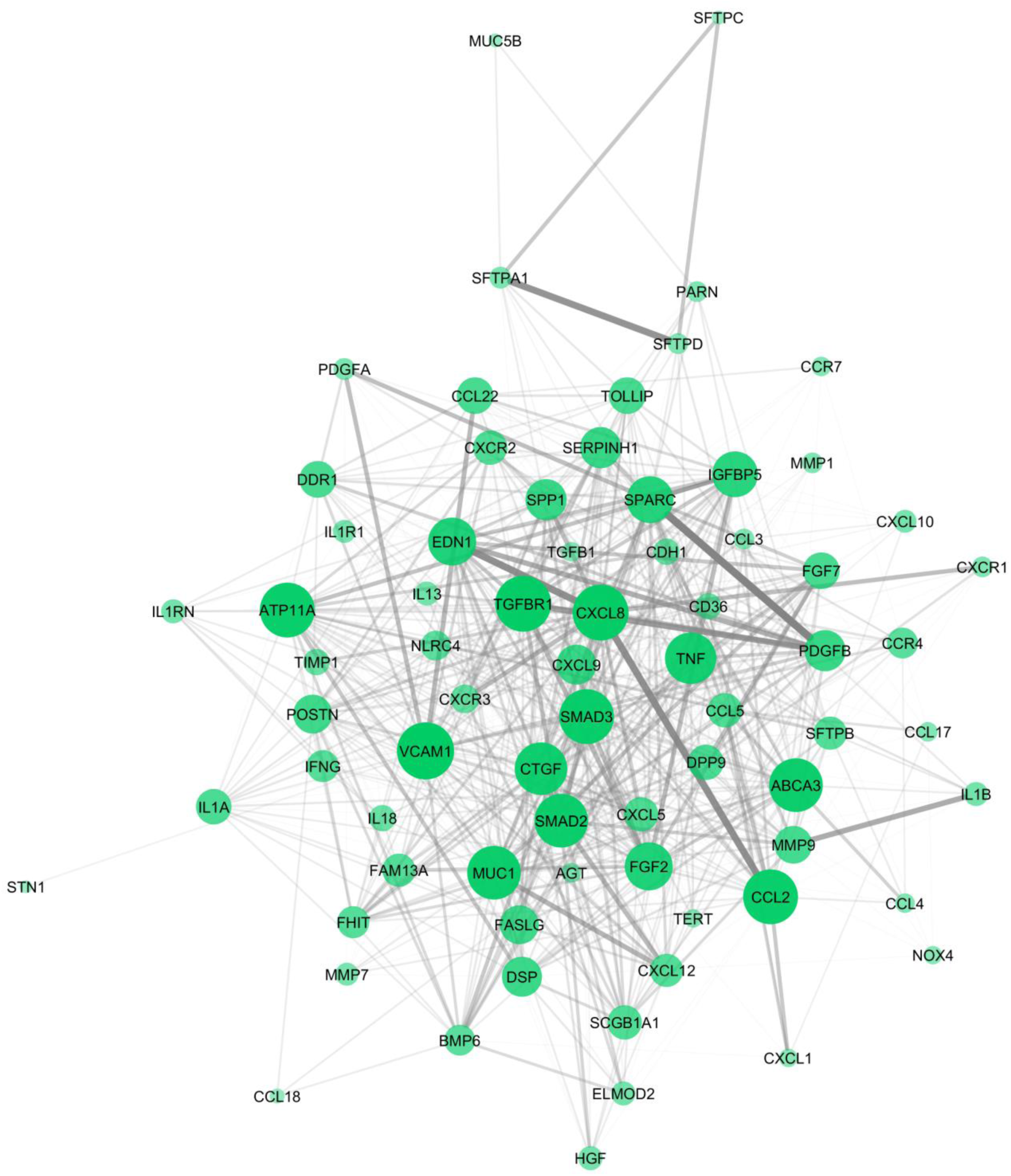
The gene map-network for the use case of IPF. The network consists of 73 gene nodes and 681 connections. The node size and transparency are based on the node’s degree and the edge width and transparency on the edge weights.

### Pathway-to-pathway Reference Network

The second component needed for the execution of PathWalks is the network of pathways, on which the walker steps, exploring pathway-to-pathway relations, in order to highlight subnetworks of disease-related molecular mechanisms. The pathways’ network is an undirected graph of functional connections that was parsed from KEGG’s KGML files. A biological pathway in KEGG consists of genes and their molecular interactions, reactions and relations. The nodes in the PathWalks’ pathway-to-pathway network represent biological pathways and an edge connecting two pathways represents a functional link between them. We are assigning a score on each edge according to the following equation:

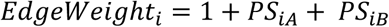

where *PS_iA_* and *PS_iB_* are the Pathway Scores (see below) of the nodes *A* and *B* connected with edge *i*.

As described in [15], the multisource integration framework combines data across various sources of information into one network and aggregates them into a gene-specific score based both on the gene characteristic information and on gene-gene integrated inter-relation. We obtain the Pathway Score (PS) of each pathway by adding the gene-specific scores of the genes participating in it. The scores of the genes carry information of the gene’s importance in the disease of study, based on the described integration method. We calculate PS scores only for the pathways that were retrieved through Enrichr’s KEGG pathway enrichment analysis [29], of the top 100 scored genes of the disease, as selected in [15].

The pathway-to-pathway networks of *AD* and *IPF* are depicted in Figures 4 and 5 respectively. In these two figures we highlight certain nodes that have high centrality scores. These pathway nodes belong to the top 5% of nodes with the highest degree and/or betweenness centralities. A node’s betweenness centrality represents the total number of the network’s shortest paths passing through the node. It is highly likely that these pathways appear in the results of PathWalks, due to the topology of the network. Similarly, thicker edges represent stronger connections based on the assigned weights and are more likely to be traversed by the walker.

**Figure 4:**
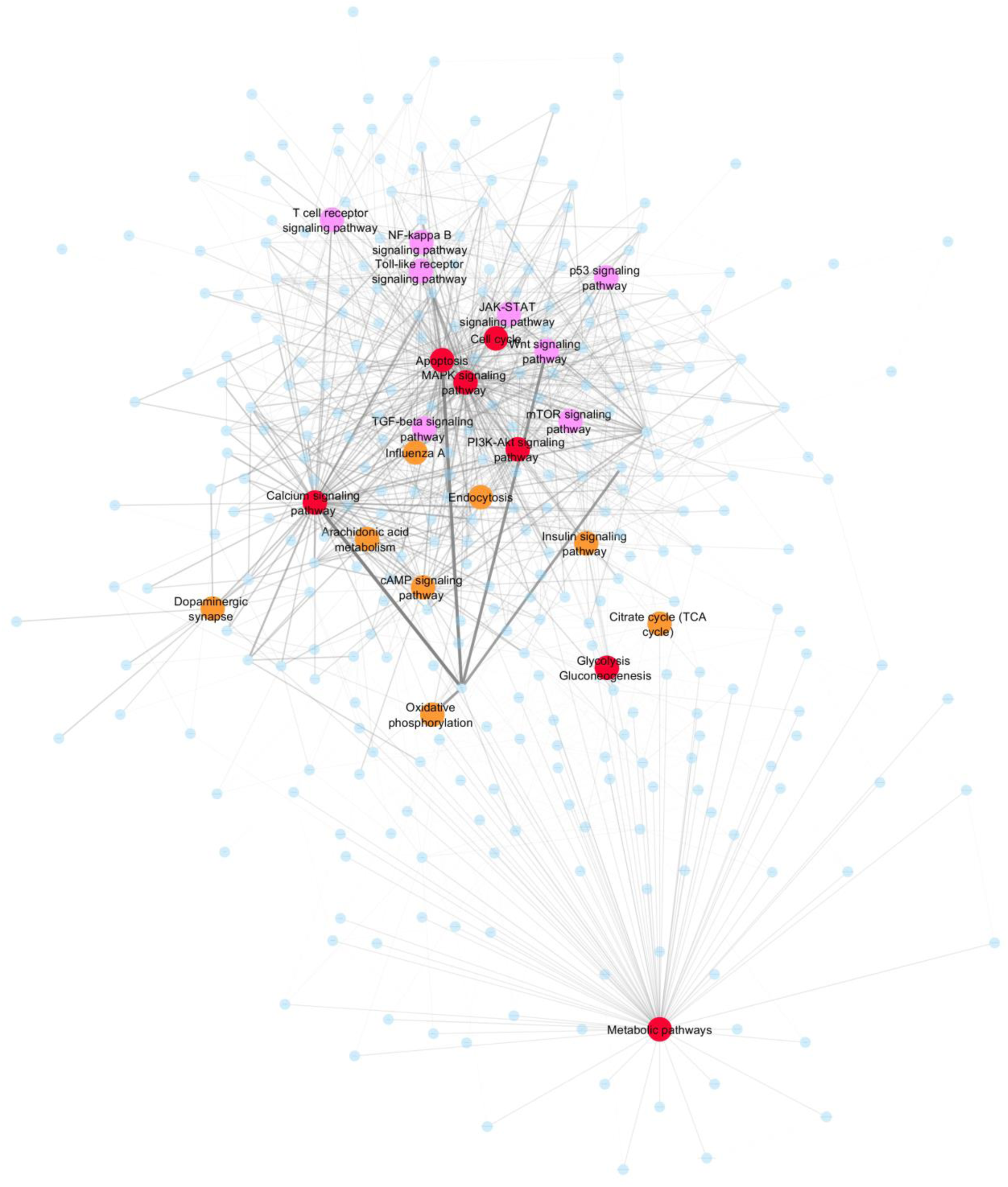
The base pathway-to-pathway network for the use case of AD. The network consists of 319 pathways and 1329 connections. The top 5% of the nodes with the highest betweenness centrality are colored orange, with the highest degree are colored pink and with both of the above are colored red. The edges’ width and transparency are relative to the starting weights of the AD pathways’ network.

**Figure 5:**
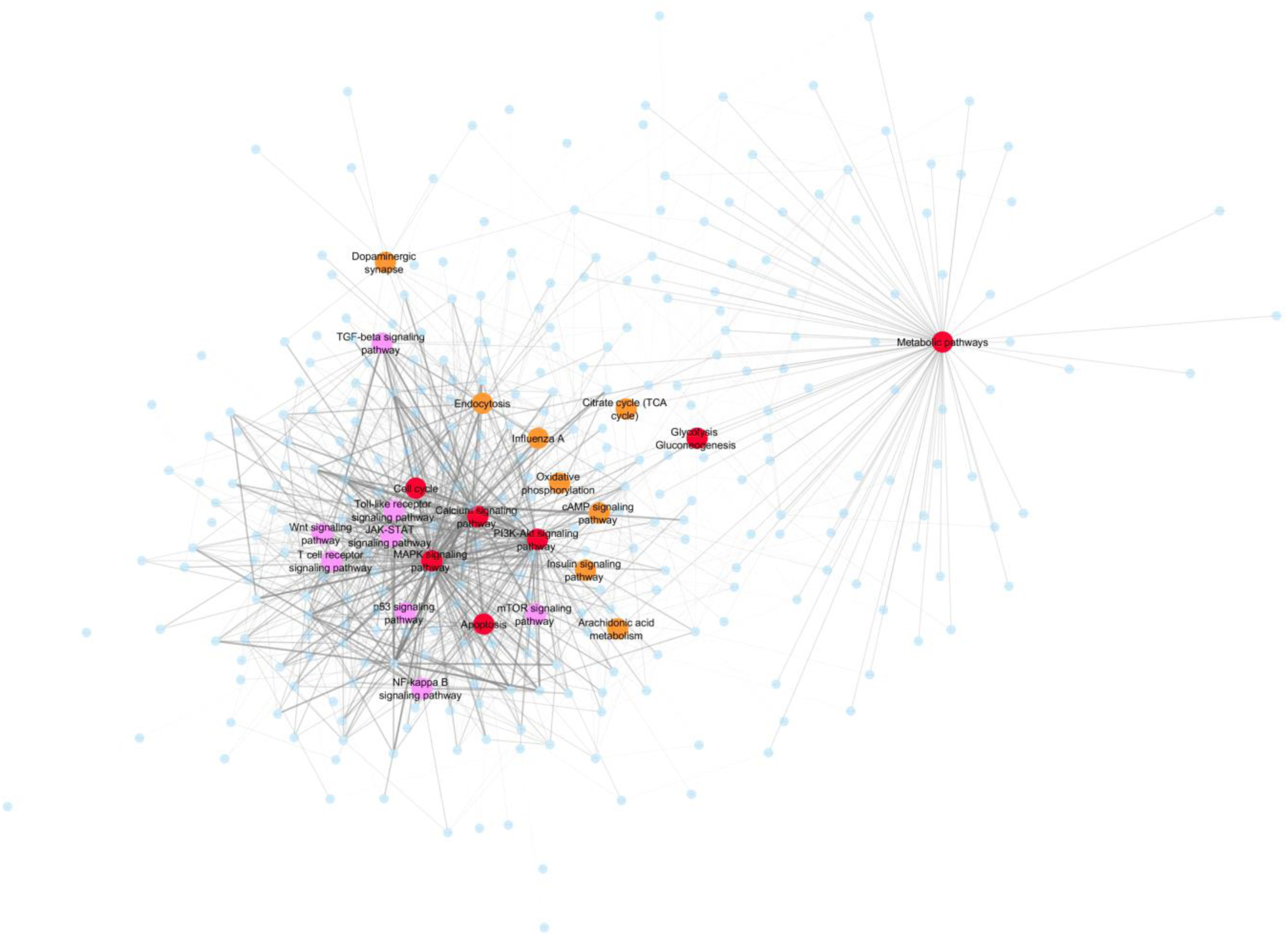
The base pathway-to-pathway network for the use case of IPF. The network consists of 319 pathways and 1329 connections. The top 5% of the nodes with the highest betweenness centrality are colored orange, with the highest degree are colored pink and with both of the above are colored red. The edges’ width and transparency are relative to the starting weights of the IPF pathways’ network.

### Pathways’ Community Detection by Accumulating Guided Tours

Following the construction of the gene map and the pathways’ network, we initiate the execution of the main algorithm. At the beginning of the execution a random gene and a random pathway starting nodes are selected, one for each of the two networks respectively (gene-gene, pathway-pathway). At each iteration, the walker performs a series of steps on the gene map level and the result assists the walker in deciding its next destination on the pathways’ level. On the genes’ network the walker moves based on a simple random walk methodology, with a random restart every fifty iterations. In more detail, a random number *n* is generated in every iteration based on a Cauchy distribution which indicates the number of steps the walker has to complete on the genes’ level. The walker traverses higher-weighted edges with higher probability via Monte Carlo sampling. The restart parameter prevents the walker from staying trapped inside neighborhoods of high-degree connectivity or bouncing between neighbors with high edge-weight values. Including the starting gene node, the maximum number of genes that can participate in a path during one turn, is *n + 1*, in the case where no nodes were visited more than once. The selected nodes indicate the next destination of the random walker on the pathways’ level.

Every pathway receives a *+1* score for each of the selected genes that is included in it. Through a second Monte Carlo sampling the next pathway is chosen, based on the traversed genes’ participation in it. Then, the walker travels the shortest path between the current and the chosen pathway node. In case of multiple shortest paths with the same score, a random one is chosen among them. If no pathways were found containing any of the traversed genes, a new random pathway is sampled and the walker travels there via the shortest path. All of the pathway nodes and all the edges participating in the selected shortest path receive a *+1* on their final score. The resulting list of the top ranked pathways highlights key molecular mechanisms, according to the genetic map of the disease of interest, while the sorted edge list result is important for the discovery of pathway communities; biological mechanisms and their relations. The results of the PathWalks algorithm tend to favor nodes with high betweenness score, due to the shortest path usage while pathway-traversing. In order to highlight the most important pathways, we pay special attention to the mostly walked pathways that do not necessarily have high betweenness values.

The PathWalks algorithm’s convergence criterion is based on the similarity index between the current and the last sorted list (every a set number steps, *100* in our use cases) of the most visited pathways. In the case that the similarity index between two pathway lists is above a defined threshold, then the walker is allowed to finish the execution. We call this defined threshold, the algorithm’s converging factor. To avoid any random high-similarity result that might occur mid-execution, the variance of the last ten similarity comparisons is calculated; if the variance is below a certain low threshold (e.g. 0.003), while at the same time the similarity index exceeds the converging factor (e.g. 95% similarity), the execution finishes. The stricter the converging factor is, the longer the algorithm needs to converge but the resulting pathway communities are less noisy and more related to the disease-related map that guided the walker on the pathway network.

We developed the PathWalks software in the R programming language [30] and used CRAN’s igraph package [31] for handling network activity. We show the pseudocode for the PathWalks algorithm in Figure 6. We also plotted all network figures using the Cytoscape tool [32].

**Figure 6:**
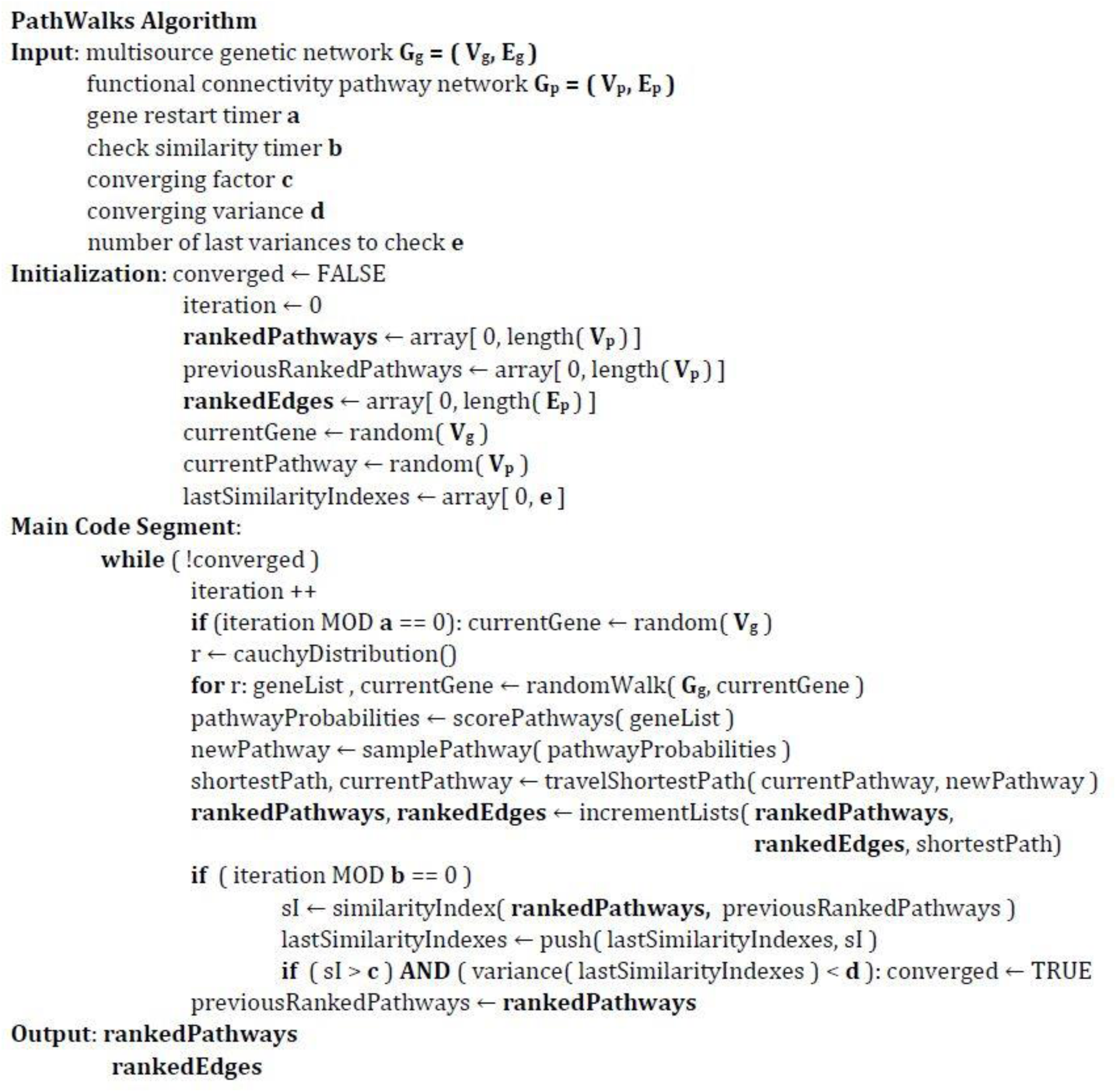
The pseudocode for PathWalks briefly describing the inputs, the execution procedure and the outputs of the algorithm.

## Case Studies’ Results

We chose *AD* and *IPF* as two use case diseases for PathWalks. Both are incurable diseases with sufficient available omics data online. Both remain active research targets and seemed appropriate for studying in order to better understand the biological mechanisms that are responsible for their pathogenesis. The PathWalks algorithm was ran iteratively until the desired converging similarity and variance output was achieved (see the methods’ section for more details). For the test runs on the two use cases, we set a converging factor of *0.95* and a converging variance of *0.003* (arbitrary values based on a number of initial trials). The similarity indexes and the respective variances are calculated every hundred steps. The diagrams of the values of the converging metrics during the execution of PathWalks for the two use-case scenarios are shown in Figure 7. The algorithm executed *50600* iterations in the use case of *AD* and *30600* for *IPF*. A faster convergence was achieved for *IPF* compared to *AD* (∼ 2/3 iterations) due to the smaller size of the guiding gene map (∼ 1/3 connections).

**Figure 7:**
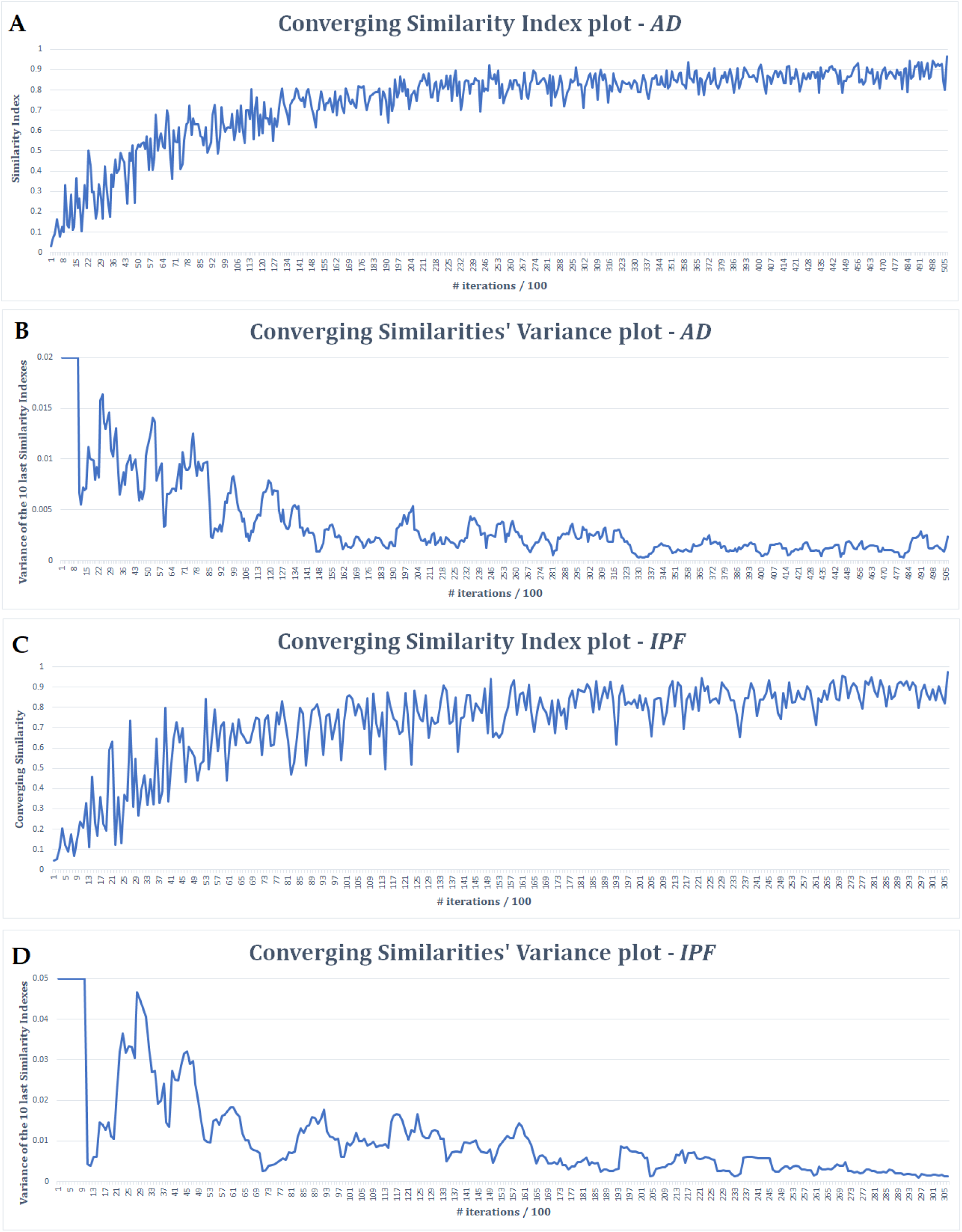
Converging similarity indexes’ plots for the AD and IPF cases. Converging similarities are calculated every hundred steps. Converging similarities’ variances are calculated for the ten last converging similarity indexes every hundred steps. A: Converging similarity index values’ plot of AD. B: Converging similarity index variance plots of AD. C: Converging similarity index values’ plot of IPF. D: Converging similarity index variance plots of IPF.

Following the convergence of PathWalks, we obtain the final resulting pathways and pathway subnetworks that are highlighted. The outputs are a list containing the most visited pathways and a list with the most frequently traversed edges. Tables 1-4 present the top ten results while Supplementary Tables 1-4 contain the ranked pathway and edge list files for the two diseases respectively. There is a certain level of self-validation in the results seeing that the “Alzheimer disease” pathway was brought to the top of the results for the use case of *AD*. Unfortunately, a respective, disease-specific pathway (such as “Pulmonary Fibrosis”) for *IPF* does not exist in the KEGG database. Figures 8 and 9 show the resulting pathway networks for the two diseases. Instead of the initial edge weights (see Figures 4 and 5) the edge thickness is now relative to the number of times the edge was walked during the algorithm and the node size is relative to the times a node was accessed. Nodes in green are brought to the top of the results according to the omics data relative to the disease (multisource map and pathway edge scores) and not necessarily due to the topology of the network, unlike the red, orange and pink nodes. Finally, Figures 10 and 11 zoom in the most frequently walked subnetwork of each disease and are created with Cytoscape’s [32] prefuse force directed layout, which brings pathway nodes, whose common edge was walked most often, closer to each other. At a glance, a key pathways’ community generated in the use case of *AD* is the “Alzheimer disease” pathway, which is strongly connected to the “Apoptosis” and “Calcium signaling pathway” pathways, and their immediate neighbors. In the use case of *IPF*, an emerging community consists of the “MAPK signaling pathway” and its closest neighbors “Pathways in cancer”, “Toll-like receptor signaling pathway”, “Chemokine signaling pathway” and “TNF signaling pathway”.

**Figure 8:**
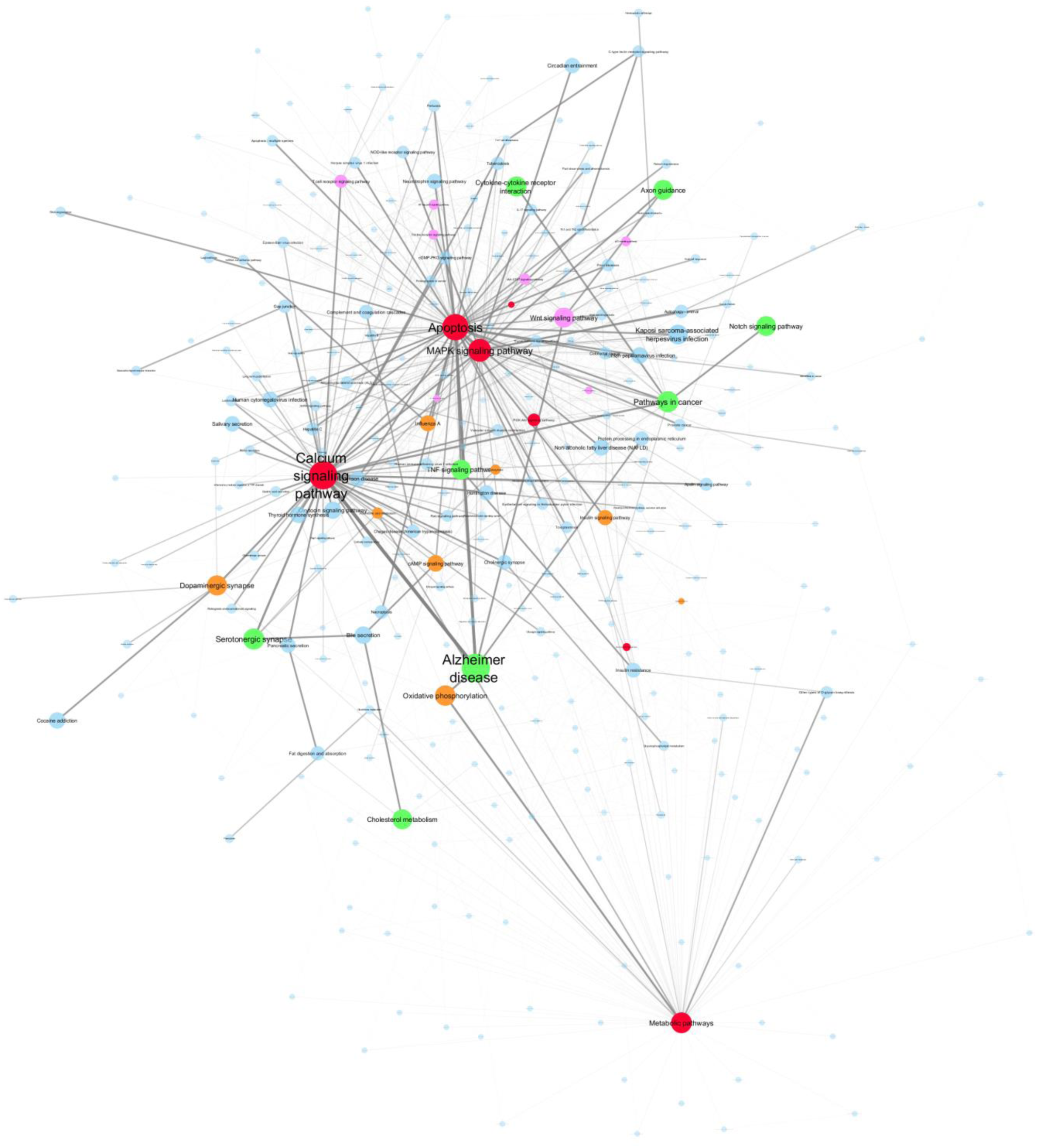
The result of the PathWalks algorithm for the AD use case. The results are overlaid over the base AD pathways’ network. The nodes’ sizes are relative to the number of times they were accessed. The top 5% walked nodes that do not belong in the starting 5% of either betweenness centrality or degree scores are colored green. The edges’ width and transparency are also relative to the times they were traversed by the walker.

**Figure 9:**
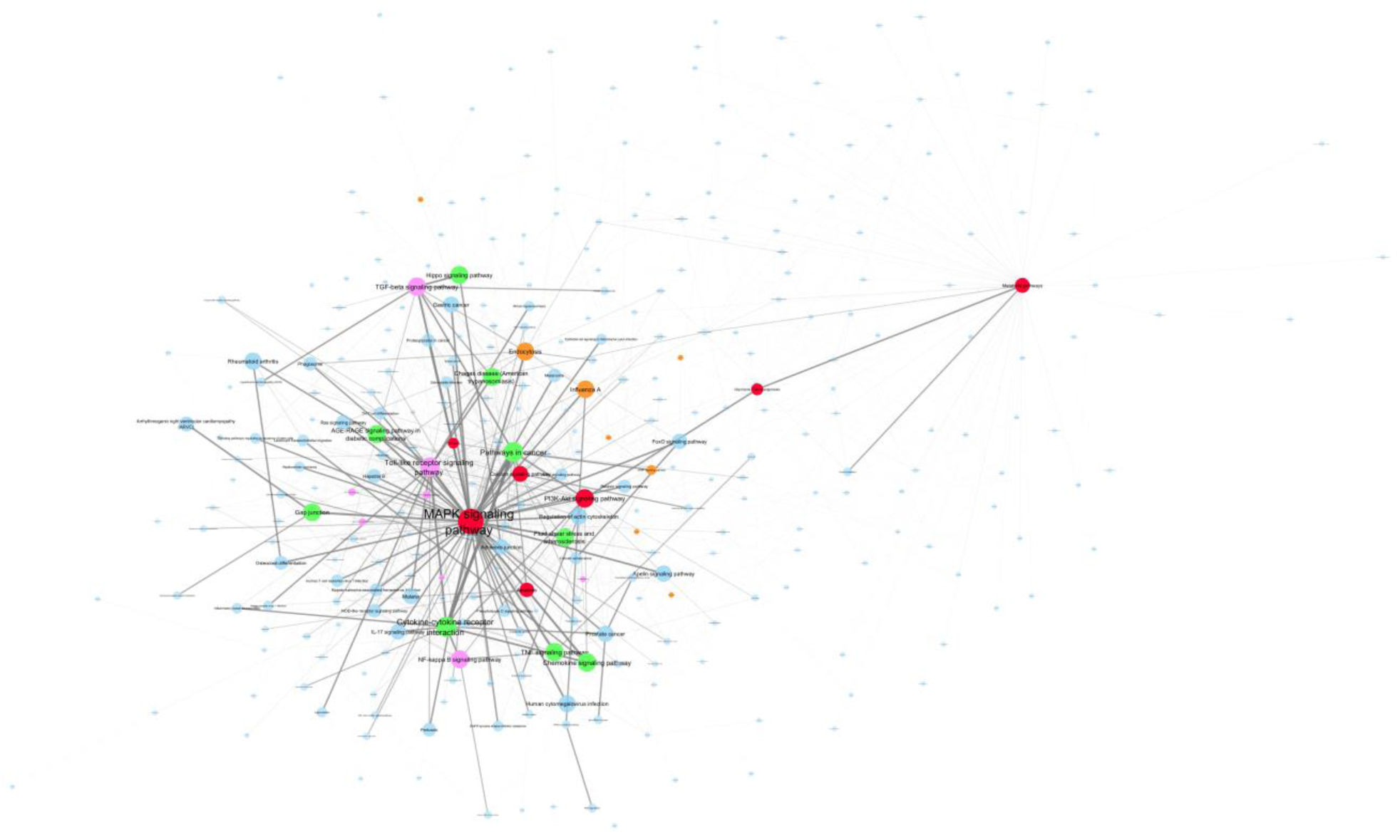
The result of the PathWalks algorithm for the IPF use case. The results are overlaid over the base IPF pathways’ network. The nodes’ sizes are relative to the number of times they were accessed. The top 5% walked nodes that do not belong to the starting 5% of either betweenness centrality or degree scores are colored green. The edges’ width and transparency are also relative to the times they were traversed by the walker.

**Figure 10:**
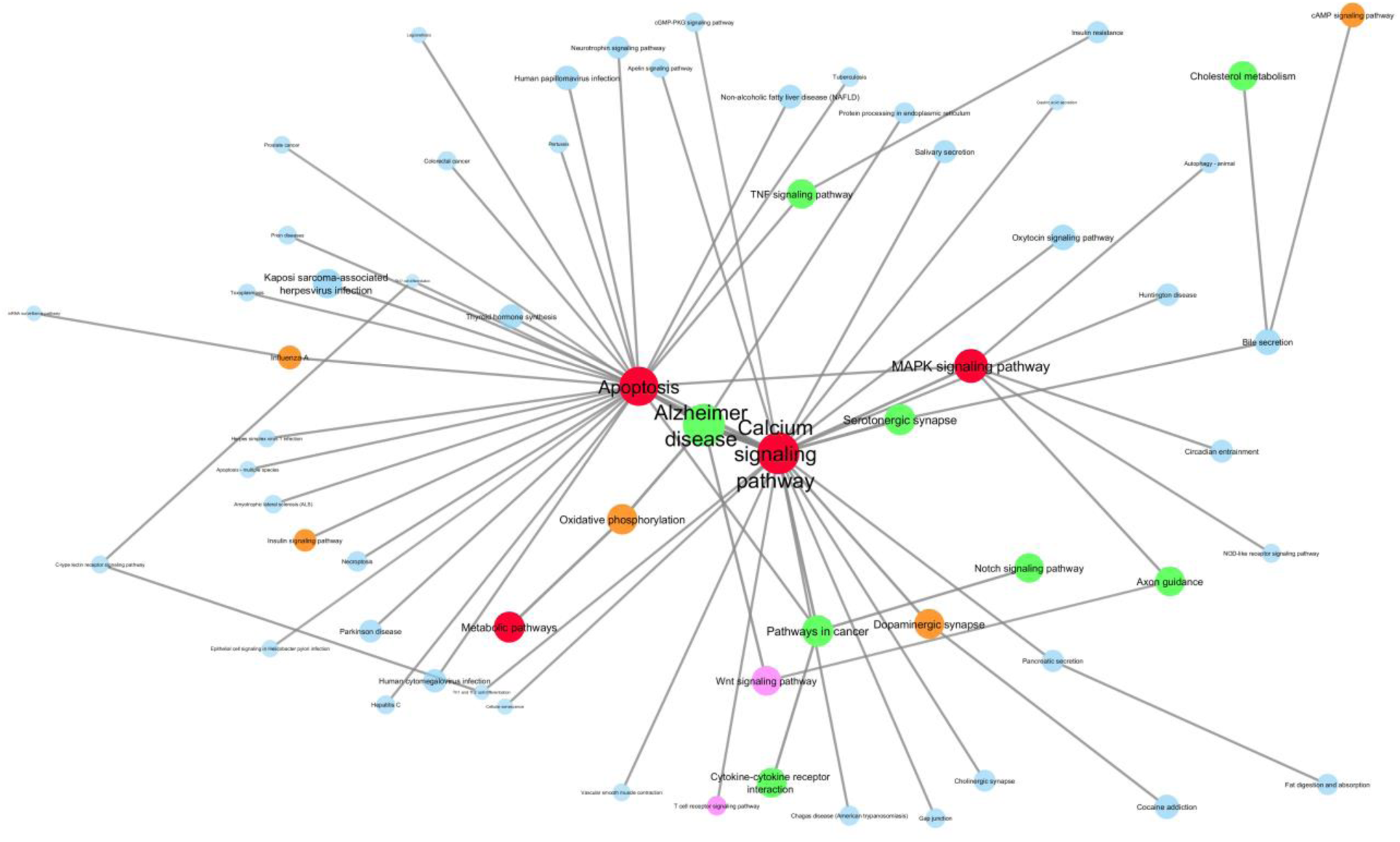
Zoom-in on the top 5% walked edges of the AD use case pathways’ network. Pathways are brought closer relatively to the number of times the edge between them was walked. A strong pathways’ community emerging from this figure is the path linking “Apoptosis” to “Alzheimer disease”, “Alzheimer disease” to “Calcium signaling pathway” and their immediate neighbors.

**Figure 11:**
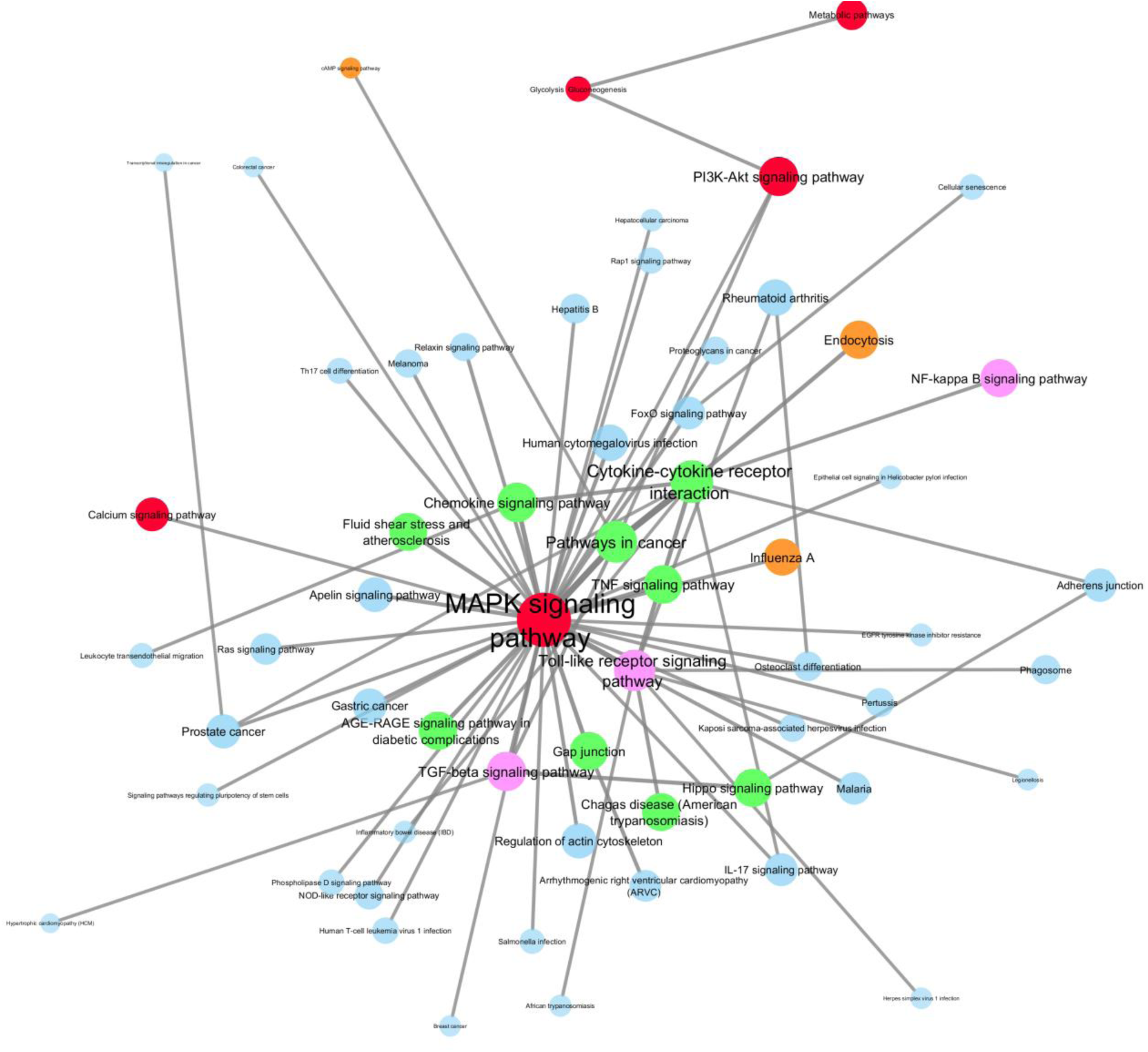
Zoom-in on the top 5% walked edges of the IPF use case pathways’ network. Pathways are brought closer relatively to the number of times the edge between them was walked. A strong pathways’ community emerging from this figure is the cluster of the “MAPK signaling pathway” and its closest neighbors “Pathways in cancer”, “Toll-like receptor signaling pathway”, “Chemokine signaling pathway” and “TNF signaling pathway”.

**Table 1:**
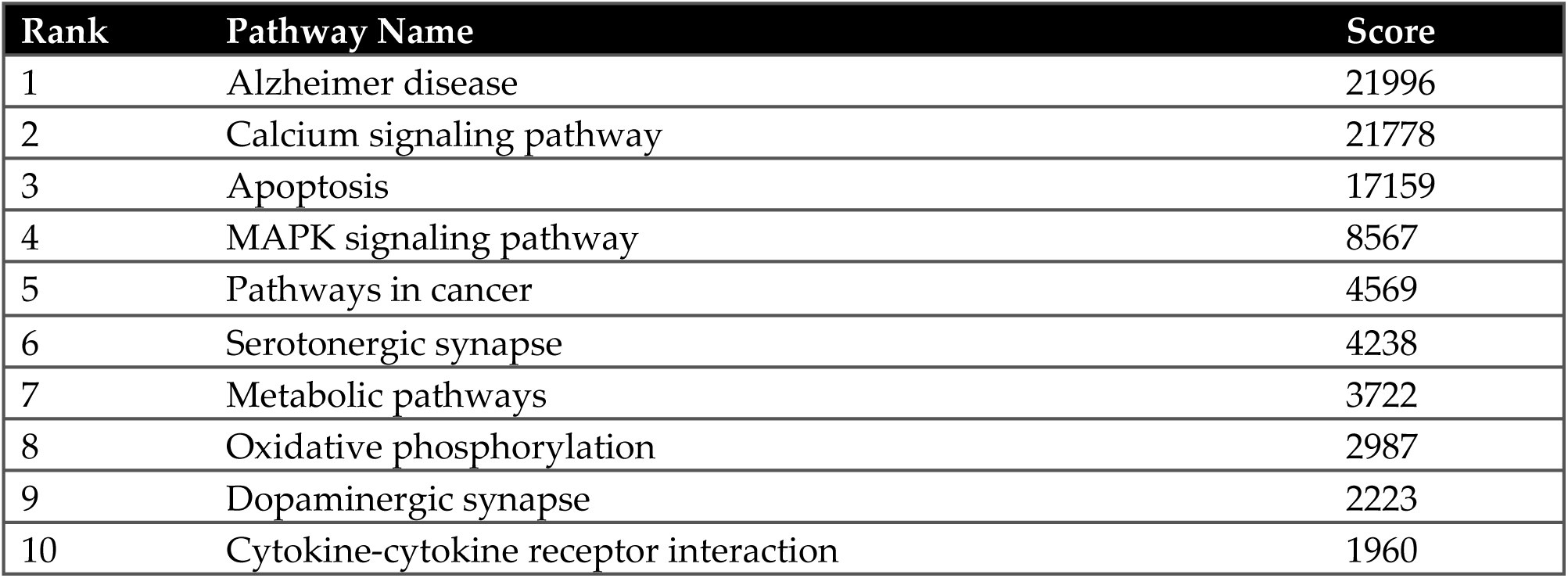
The top-ten ranked pathways that are visited in the use case of *AD*. The score denotes the times a pathway participated in the shortest path that was traversed by the random walker.

**Table 2:** The top ten ranked edges walked in the use case of *AD*. The edge weight denotes the number of times an edge was accessed by the random walker.

| Rank | Pathway Name 1 | Pathway Name 2 | Edge Weight |
| --- | --- | --- | --- |
| 1 | Alzheimer disease | Calcium signaling pathway | 6427 |
| 2 | Alzheimer disease | Apoptosis | 4736 |
| 3 | Calcium signaling pathway | Serotonergic synapse | 1905 |
| 4 | Oxidative phosphorylation | Alzheimer disease | 1410 |
| 5 | Oxidative phosphorylation | Metabolic pathways | 1389 |
| 6 | Pathways in cancer | Calcium signaling pathway | 1268 |
| 7 | MAPK signaling pathway | Calcium signaling pathway | 1204 |
| 8 | Calcium signaling pathway | Dopaminergic synapse | 1015 |
| 9 | Pathways in cancer | Cytokine-cytokine receptor interaction | 841 |
| 10 | Alzheimer disease | Wnt signaling pathway | 776 |

**Table 3:** The top-ten ranked pathways that are visited in the use case of *IPF*. The score denotes the times a pathway participated in the shortest path that was traversed by the random walker.

| Rank | Pathway Name | Score |
| --- | --- | --- |
| 1 | MAPK signaling pathway | 17942 |
| 2 | Cytokine-cytokine receptor interaction | 6105 |
| 3 | Pathways in cancer | 5263 |
| 4 | Toll-like receptor signaling pathway | 4598 |
| 5 | Chemokine signaling pathway | 2709 |
| 6 | TGF-beta signaling pathway | 2566 |
| 7 | PI3K-Akt signaling pathway | 2434 |
| 8 | TNF signaling pathway | 1783 |
| 9 | Endocytosis | 1481 |
| 10 | AGE-RAGE signaling pathway in diabetic complications | 1469 |

**Table 4:** The top ten ranked edges walked in the use case of *IPF*. The edge weight denotes the number of times an edge was accessed by the random walker.

| Rank | Pathway Name 1 | Pathway Name 2 | Edge Weight |
| --- | --- | --- | --- |
| 1 | Pathways in cancer | Cytokine-cytokine receptor interaction | 1430 |
| 2 | Pathways in cancer | MAPK signaling pathway | 1393 |
| 3 | MAPK signaling pathway | Toll-like receptor signaling pathway | 941 |
| 4 | MAPK signaling pathway | Chemokine signaling pathway | 722 |
| 5 | MAPK signaling pathway | TNF signaling pathway | 669 |
| 6 | Cytokine-cytokine receptor interaction | Toll-like receptor signaling pathway | 610 |
| 7 | MAPK signaling pathway | TGF-beta signaling pathway | 606 |
| 8 | MAPK signaling pathway | Gap junction | 599 |
| 9 | MAPK signaling pathway | AGE-RAGE signaling pathway in diabetic complications | 589 |
| 10 | Cytokine-cytokine receptor interaction | Endocytosis | 549 |

## Discussion

The PathWalks methodology combines random walks and network-based integration to detect key disease-related pathway communities. In this paper, we apply the PathWalks methodology on the use cases of *AD* and *IPF* for pathway community detection. The most visited pathway in the use case of *AD* is, as expected, the “Alzheimer disease” pathway which includes a set of known components and interactions related to the *AD* pathology. The second most visited pathway is the “Calcium signaling pathway” which has the strongest connection to the “Alzheimer disease” pathway based on the most walked edges of the pathways network. Calcium dysfunction has been associated with several neurodegenerative diseases including *AD* [33–35]. Alteration in calcium homeostasis has been found in *AD* animal models, leading to elevated levels of resting calcium [36]. Calcium overload has also been correlated with disrupted neuronal structure and function [37].

The “Alzheimer disease” pathway is also linked indirectly through the “Calcium signaling pathway” with frequently walked edges to other pathways such as the “Serotonergic synapse”, “Dopaminergic Synapse” and “MAPK signaling pathway”. The persistent activation of mitogen-activated protein kinases (MAPKs) is thought to play a key role in neurodegeneration, including *AD*, through mediating hyper-phosphorylation of neuronal proteins, eventually causing neuronal death [38]. Note that the “Apoptosis” pathway is highlighted as the second most visited edge in the *AD* network linked to the “Alzheimer disease” pathway. The direct and indirect relationship of MAPK signaling cascades and calcium is also well documented [39]. The serotonergic system has been found to be impaired in *AD* where extensive serotonergic denervation is observed [40]. A deficit in the dopaminergic system has also been observed in *AD*, with the loss of that dopaminergic neurons in the ventral tegmental area (VTA) during the early (pre-plaque) stages of AD [41]. Both dopaminergic and serotonergic can be associated to *AD* through the calcium pathway. For example, a T-type calcium channel enhancer (known as SAK3) was shown to enhance serotonin and dopamine and releases in the hippocampus of both naive and amyloid precursor protein knock-in mice [42].

“Cytokine-cytokine receptor interaction” and “Pathways in Cancer” are two linked highlighted pathways associated with *AD* and connect to the “Calcium signaling pathway”. Interestingly certain types of cancers, such as lung cancer, have been found to be anti-correlated with the occurrence of neurodegenerative diseases such as *AD*, although both types of diseases are associated to aging [43]. Neuroinflammation is a prominent feature of *AD* in which the over-activation microglial results in an increased production of pro-inflammatory cytokines [44]. Cytokines also have an important role in cancer as they can, in certain cases, inhibit and in others facilitate cancer progression [45]. We also observe a branch of the graph connecting “Metabolism” to “Alzheimer disease” pathways, through the “Oxidative phosphorylation”, with highly scored edges. Both the hypometabolism and oxidative stress have been implicated as key contributors in initiation and progression for the synapse vulnerability in *AD* [46]. The link from “Alzheimer disease” to “Wnt signaling” is also highlighted in our results. Deregulation of the Wnt signaling pathway has been observed both in the aging brain as well as in *AD* with implications for the relevant synaptic pathology [47]. Nevertheless, “Metabolism”, “Oxidative phosphorylation” and “Wnt signaling” are pathways with high betweenness and degree scores and probably act as intermediate functional nodes in *AD*.

For the case of *AD*, in the top 20 visited pathways, the “Kaposi sarcoma-associated herpesvirus infection” (HHV-8) and the “Human papillomavirus infection” (HPV) pathways are also found. HPV is thought to be responsible for more than 90% of anal and cervical cancers and about 40% of vaginal, vulvar and penile cancers. Infection with high-risk human papillomaviruses (HPVs) has also recently been implicated in the pathogenesis of head and neck squamous cell carcinomas (HNSCCs) [48]. Kaposi sarcoma is caused by infection with a virus called the Kaposi sarcoma associated herpesvirus (KSHV), also known as human herpesvirus 8 (HHV8). KSHV is in the same family as Epstein-Barr virus (EBV), the virus that causes infectious mononucleosis (mono) and is linked to several types of cancer [49]. Recent studies have found a strong link between the activity of specific viral species (three herpesviruses: HSV1 HHV-6 and HHV-7), with molecular, genetic, clinical, and neuropathological aspects of *AD* [50, 51]. It is worth noting that in our analysis “Kaposi sarcoma-associated herpesvirus infection” and “Human papillomavirus infection” pathways are also indirectly linked with the “Alzheimer disease” pathway through the “Apoptosis” pathway (top 20 walked edges of *AD*). As pathogens, especially viruses, are increasingly involved in diseases with no apparent association, it is worth further investigating their involvement in the development and progression of *AD*.

In the use case of *IPF* we find the “MAPK signaling pathway” to be ranked on top, based on the walker’s visitation frequency. “MAPK signaling pathway” has high betweenness and degree scores, but is linked to other highlighted pathways of *IPF* and hence might be a key intermediate functional node in the pathogenesis of *IPF*. Antoniou et al. [52] observed a significant overexpression in the Braf oncogene, a key gene in the MAPK pathway, in an *IPF* vs a control group study. In another study, Yoshida et al. [53] suggested three MAP kinases (ERK, JNK and p38 MAPK) to be involved in the regulation of lung inflammation and injury in *IPF*. Additionally, we have suggested in our previous computational drug repurposing study on *IPF* [54] that the MAPK signaling pathway seems to play a key role in the transition of early stage *IPF* towards a more advanced stage.

The second highest ranked pathway in *IPF* is “Cytokine-cytokine receptor interaction”. This has been suggested by our previous study [54] to be a key pathway in all stages of the *IPF* disease. The important role of cytokines as therapeutic targets in *IPF* has also been emphasized by Coker et al. [55]. Bouros et al. recently proposed the Tumor necrosis factor-Like cytokine 1 A (TL1A), as a novel fibrogenic factor [56]. Specifically, they found upregulated mRNA and protein levels of TL1A in subepithelial lung myofibroblasts (SELMs) that were treated either with pro-inflammatory factors or bronchoalveolar lavage fluid (BALF) from *IPF* patients.

“Pathways in cancer” and “Toll-like receptor signaling pathway” are the next two mechanisms in rank. *IPF* has been known to have many similar alterations and behaviors to cancer biology [57]. The two most walked edges link the “Cytokine-cytokine receptor interaction” pathway to the “Pathways in cancer” which is then linked to the “MAPK signaling pathway”. Yong et al. [58] presented information about p38 MAPK being a key player in cellular processes that are related to inflammation and cancer. p38 MAPK can activate both anti-inflammatory and pro-inflammatory cytokines. p38 MAPK inhibitors have been tested as potential therapeutic drugs against inflammatory diseases and cancer but with numerous side effects.

Recent Toll-like receptor studies related to *IPF* suggest promising genes as therapeutic targets. TLR7, TLR9 and TLR2 mRNA expressions were found to be significantly increased in *IPF* compared to control subjects, even though TLR9 protein expression was lower in *IPF* than controls [59]. TLR9 has also been shown to drive the fibrosis progression in *IPF* in the study of Hogaboam et al. [60]. A TLR3 polymorphism, namely TLR3 L412F, has also been linked to a more aggressive and profibrotic disease phenotype in *IPF* [61]. The third most walked edge links “MAPK signaling pathway” and “Toll-like receptor signaling pathway”. In a regulatory network the edge would be directed from the “Toll-like receptor signaling pathway” towards the MAPK one, since TLR signaling leads to the activation of MAPKs in mammals through the sequential recruitment of the adapter molecule MyD88 and the serine-threonine kinase IRAK [62]. In turn, the activated MAPKs (ERKs, JNKs and p38 proteins) regulate cellular mechanisms associated with inflammatory responses as well as cell proliferation and survival [63], which are key components in the pathogenesis of *IPF*.

The “Chemokine signaling pathway” (fifth in rank) has also been shown to contribute to the pathogenesis of interstitial lung diseases like *IPF* via mechanisms such as the regulation of vascular modeling and the mediation of the traffic of bone marrow derived progenitor cells to the lungs [64]. The fourth most-walked edge links “Chemokine signaling pathway” to “MAPK signaling pathway”. Interleukin-8 (IL8 or CXCL8) is a chemokine known to activate MAPK through the activation of RAF1 and BRAF genes [65]. The “TGF-beta signaling pathway” (sixth in rank) is also known to be linked not only with *IPF* but with fibrotic diseases in general [66] and it is known to be one of the key drivers in fibrogenesis [67].

In the top 40 visited pathways of *IPF*, the “NOD-like receptor signaling pathway” is also found. A relationship between total bacterial load and poor prognosis in *IPF* samples has been reported. More specifically, an enriched detection of organisms such as Haemophilus, Neisseria Streptococcus and Veillonella has been found in the BAL fluid of *IPF* patients [68]. Moreover, it has been observed that augmented concentrations of certain strains of Streptococcus, in BAL samples from *IPF* patients, associate with increased NOD receptor signaling and poor outcomes [69]. The potential contribution of bacteria in *IPF* pathogenesis is also an attractive area of investigation for novel treatment approaches [70]. It should be mentioned that, in our analysis, “NOD-like receptor signaling pathway” is linked with “MAPK signaling pathway”, a central molecular pathway of *IPF*.

Without doubt, there is no ground truth in order to validate the highlighted pathway communities apart from looking into the literature or carrying out lab experiments. However, conceptually, this work gives meaningful results regarding the disease of interest given that the pathway-to-pathway network and the gene-map are carrying biological information.

## Acknowledgments

Evangelos S. Karatzas is a PhD student in the National and Kapodistrian University of Athens. His doctoral thesis is being funded by the IKY (State Scholarships Foundation) scholarship, funded by the Action “Strengthening Human Resources, Education and Lifelong Learning”, 2014-2020, co-funded by the European Social Fund (ESF) and the Greek State. Marilena M. Bourdakou is a Postdoctoral Researcher in the Democritus University of Thrace. Her postdoctoral research is being funded by the IKY (State Scholarships Foundation) scholarship, funded by the Action “Strengthening Human Resources, Education and Lifelong Learning”, 2020-2022, co-funded by the European Social Fund (ESF) and the Greek State. Margarita Zachariou, George Minadakis and Anastasios Oulas hold a postdoctoral research fellow position funded by the European Commission Research Executive Agency Grant BIORISE (No. 669026), under the Spreading Excellence, Widening Participation, Science with and for Society Framework. George M. Spyrou holds the Bioinformatics ERA Chair Position funded by the European Commission Research Executive Agency (REA) Grant BIORISE (Num. 669026), under the Spreading Excellence, Widening Participation, Science with and for Society Framework.

